# Defining the signalling determinants of a posterior ventral spinal cord identity in human neuromesodermal progenitor derivatives

**DOI:** 10.1101/2020.06.24.168625

**Authors:** Matthew Wind, Antigoni Gogolou, Ichcha Manipur, Ilaria Granata, Larissa Butler, Peter W. Andrews, Ivana Barbaric, Ke Ning, Mario R. Guarracino, Marysia Placzek, Anestis Tsakiridis

## Abstract

The anteroposterior axial identity of motor neurons (MNs) determines their functionality and vulnerability to neurodegeneration. Thus it is a critical parameter in the design of strategies aiming to produce MNs from human pluripotent stem cells (hPSCs) for regenerative medicine and disease modelling applications. However, the *in vitro* generation of posterior spinal cord MNs has been challenging. Although the induction of cells resembling neuromesodermal progenitors (NMPs), the *bona fide* precursors of the mammalian spinal cord, offers a promising solution, the progressive specification of posterior MNs from these cells is not well-defined. Here we determine the signals guiding the transition of human NMP-like cells toward posterior ventral spinal cord neurectoderm. We show that combined WNT-FGF activities drive a posterior dorsal early neural state while suppression of TGFβ-BMP signalling pathways, combined with SHH stimulation, promotes a ventral identity. Based on these results, we define an optimised protocol for the generation of posterior MNs that can efficiently integrate within the neural tube of chick embryos. We expect that our findings will facilitate the functional comparison of hPSC-derived spinal cord cells of distinct axial identities.

## Introduction

The developing spinal cord generates a diverse range of cell types including motor neurons (MNs). These control all muscle movements in the body and are therefore of great clinical importance in neurodegenerative conditions such as amyotrophic lateral sclerosis (ALS), as well as traumatic spinal cord injuries that cause paralysis. An attractive platform for the *in vitro* modelling and treatment of these conditions involves the generation of MNs from human pluripotent stem cells (hPSCs) via directed differentiation or transcriptional programming approaches (Davis-Dusenbery et al., 2014). During embryonic development, MNs arise from ventrally located spinal cord progenitors and subsequently become organised into longitudinally arrayed columns and pools connected to various muscle targets (reviewed in Sagner and Briscoe, 2019). The induction of MN subtypes and their subsequent muscle target specificity is determined by their position along the anteroposterior (A-P) axis. The A-P position of MNs appears to also influence their vulnerability to ALS (Brockington et al., 2013; Gerardo-Nava et al., 2013; Kaplan et al., 2014). Thus the *in vitro* production of spinal cord cells/MNs of defined A-P axial identities from hPSCs is a critical factor to consider in the design of regenerative medicine therapies and the study of axial level-specific neuroprotective mechanisms.

The A-P patterning of spinal cord MNs is orchestrated primarily by the coordinated action of *Hox* gene family members (Dasen et al., 2008; Dasen et al., 2003; Dasen et al., 2005; Jung et al., 2010; Lacombe et al., 2013; Mendelsohn et al., 2017; Philippidou and Dasen, 2013). *Hox* genes are arranged as paralogous groups (PG) (1 to 13) across four distinct chromosomal clusters (A, B, C and D) and they are expressed along the post-cranial A-P axis in a strict spatiotemporal manner reflecting their 3’-5’ genomic order (reviewed in (Philippidou and Dasen, 2013)). Hindbrain/cervical spinal cord MNs innervating facial, neck/shoulder and diaphragm muscles are marked by the expression of *Hox* PG(1-5) members. Posterior brachial/thoracic spinal cord MNs innervating the upper limbs, hypaxial muscles and sympathetic ganglia are characterised by *Hox* PG(6-9) expression. MNs at the lumbosacral axial levels controlling lower limb movements are defined by expression of *Hox* PG(10-13) members. The preservation of this expression pattern in the adult spinal cord suggests additional role(s) for *Hox* genes in controlling axial identity–dependent cell function in the spinal cord after birth (Nichterwitz et al., 2016; Sabourin et al., 2009; Zeisel et al., 2018).

To date, most protocols aiming to generate ventral spinal cord cells and MNs from hPSCs induce predominantly cells of a mixed hindbrain/cervical axial identity marked by expression of *Hox* PG(1-5) members but are inefficient in producing high numbers of more posterior, thoracic/lumbosacral *Hox* PG(6-13)-positive spinal cord cells. The advent of culture regimens (both adherent and 3-dimensional) promoting the induction of neuromesodermal progenitor (NMP)-like cells *in vitro* has opened new promising avenues toward the *in vitro* generation of “hard-to-make” posterior MNs (Faustino Martins et al., 2020; Gouti et al., 2014; Kumamaru et al., 2018; Libby, 2020; Lippmann et al., 2015; Rayon, 2019; Turner et al., 2014; Verrier et al., 2018). NMPs are the posteriorly-located multipotent cell population that gives rise to both ventral spinal cord and paraxial mesoderm/somites in amniote embryos (Cambray and Wilson, 2007; Guillot, 2020; Mugele, 2018; Tzouanacou et al., 2009; Wood, 2019; Wymeersch et al., 2016 ; reviewed in Henrique et al., 2015) and is marked by the co-expression of early neural and mesodermal genes such as *Sox2* and *Brachyury (T)* (Olivera-Martinez et al., 2012; Tsakiridis et al., 2014; Wymeersch et al., 2016). However, the yields of posterior MN populations produced via NMP-based protocols have been either undefined or low, especially in the case of 2-dimensional, adherent differentiation protocols (Estevez-Silva et al., 2018; Lippmann et al., 2015). This suggests that thorough characterisation of the progressive lineage restriction events underlying the neural differentiation of human NMPs *in vitro* is required for the optimisation of current posterior MN differentiation protocols.

*In vivo*, the induction/maintenance of NMPs is directed by the WNT and FGF signalling pathways (Amin et al., 2016; Anderson, 2020; Boulet and Capecchi, 2012; Cunningham et al., 2015; Delfino-Machin et al., 2005; Diez del Corral et al., 2002; Diez del Corral et al., 2003; Garriock et al., 2015; Goto et al., 2017; Jurberg et al., 2014; Nordstrom et al., 2006; Olivera-Martinez and Storey, 2007; Osorno et al., 2012; Takemoto et al., 2006; Wymeersch et al., 2016; Young et al., 2009), which also drive the production of NMPs from PSCs *in vitro* (Gouti et al., 2014; Lippmann et al., 2015; Turner et al., 2014). Downstream neural differentiation of embryonic NMPs relies on high levels of somite-derived retinoic acid (RA) signalling acting on an FGF-dependent pre-neural/early neural precursor (Diez del Corral et al., 2002; Diez del Corral et al., 2003), while high WNT and FGF activities drive paraxial mesoderm specification (Garriock et al., 2015; Goto et al., 2017; Takada et al., 1994; Takemoto et al., 2011). WNT signals also act on *Cdx* transcription factors to trigger the acquisition of a posterior axial identity in NMPs and their derivatives (Amin et al., 2016; Delfino-Machin et al., 2005; Metzis et al., 2018; Nordstrom et al., 2006; van den Akker et al., 2002; Wymeersch et al., 2019; Young et al., 2009 ; reviewed in Deschamps and Duboule, 2017; Deschamps and van Nes, 2005). Moreover, dorsoventral (D-V) patterning of NMP-derived neurectoderm is mediated by the antagonistic action of pro-dorsal WNT/BMP and pro-ventral sonic hedgehog (SHH) signals (reviewed in Placzek and Briscoe, 2018; Sagner and Briscoe, 2019).

Here we examine in detail the signal combinations directing the stepwise induction of posterior and ventral identities during the differentiation of adherent hPSC-derived NMP-like cells. We show that sustained WNT and high FGF signalling activity is sufficient to drive the transition of TBXT^+^(the human homologue of Brachyury)-SOX2^+^ cells toward a pre-neural/early neurectodermal state in the absence of exogenous RA supplementation. This transition is accompanied by upregulation of *HOX* genes, a hallmark of posterior axial identity acquisition. WNT and FGF signals also promote a default dorsal identity and we demonstrate that efficient ventralisation of WNT/FGF-induced neurectoderm requires active suppression of the pro-dorsal TGFβ and BMP signalling pathways in addition to SHH signalling stimulation. Based on these findings, we define an optimised protocol for the generation of ventral spinal cord progenitors and MNs of a thoracic axial identity and show that the resulting cells can efficiently integrate and establish appropriate neuronal projections within the posterior neural tube of chick embryos following transplantation. We anticipate that our findings will pave the way for the development of novel cell replacement strategies aiming to treat posterior spinal cord injuries and enable the mechanistic dissection of the effect of axial identity on MN vulnerability to ALS.

## Results

### WNT and FGF induce posterior dorsal neurectoderm from human NMPs

The role of WNT and FGF signalling in NMP ontogeny *in vitro* has been unclear. Previous studies showed that their combined stimulation drives the induction of NMP-like cells from PSCs in the short term (Denham et al., 2015; Frith et al., 2018; Gouti et al., 2014; Lippmann et al., 2015) as well as their differentiation toward both neurectoderm and paraxial mesoderm following longer treatment (Diaz-Cuadros et al., 2020; Edri et al., 2019; Frith et al., 2018; Gouti et al., 2014; Tsakiridis and Wilson, 2015). We thus sought to test whether different WNT/FGF levels drive distinct cell fate decisions in human NMP-like cells. TBXT^+^SOX2^+^ cultures were generated following a 3-day treatment of hPSCs with recombinant FGF2 (20 ng/ml) and the WNT agonist/GSK-3 inhibitor CHIR99021 (CHIR) (3 µM) as described previously (Frith et al., 2018; Frith and Tsakiridis, 2019). NMP-like cells were subsequently re-plated and cultured in the presence of various different amounts of FGF2-CHIR and for distinct periods (**Fig. 1A**). We found that high levels of CHIR (8 µM) and FGF2 (40 ng/ml) promoted the generation of presomitic/paraxial mesoderm cells, marked by upregulation of *TBXT, MSGN1* and *PAX3* and downregulation of *SOX2* over a 2-day period (Days 3-5) (**Fig. 1Aiii, S1A, S1B**). In contrast, longer 4-day exposure of re-plated NMP-like cells in the presence of the same WNT/FGF agonist amounts that promoted their initial induction (3 µM CHIR and 20 ng/ml FGF2), resulted in progressive reduction of TBXT-expressing cells/TBXT protein levels (**Fig. 1Ai, 1B)**, which was further enhanced when higher levels of FGF2 were employed (100 ng/ml) in combination with CHIR (**Fig. 1AiI, 1B, S1C)**. These data indicate that distinct levels and durations of WNT and FGF stimulation trigger different cell fate decisions in hPSC-derived NMP-like cells: high WNT activity combined with FGF signalling promotes the generation of paraxial mesoderm whereas lower WNT and high FGF levels drive the extinction of TBXT^+^SOX2^+^ progenitors and limit mesodermal differentiation.

**Figure 1.**
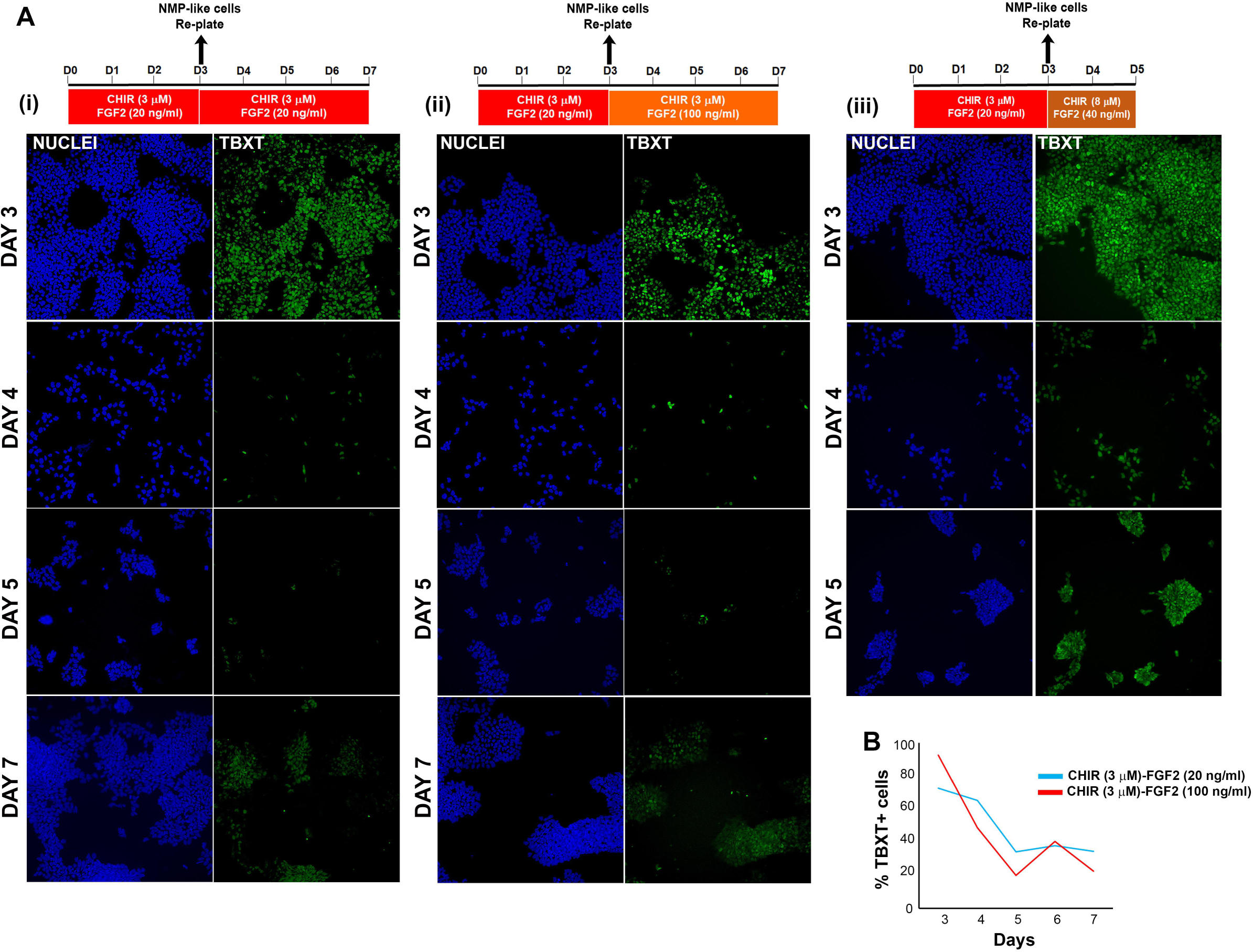
Effect of distinct levels of WNT and FGF on NMP-like cells. **(A)** Time-course immunofluorescence analysis of the expression of TBXT protein in NMP-like cells (Day 3) and their differentiating derivatives following dissociation, re-plating and culture using the indicated combinations of WNT and FGF agonists. **(B)** Quantification of the percentage of TBXT^+^ cells at different time points during treatment of NMP-like cells with the indicated combinations of WNT and FGF agonists.

We next tested whether the dramatic TBXT reduction in day (D) 7 CHIR (3 µM) – FGF2 (100 ng/ml)-treated cultures is a consequence of the acquisition of a neural identity. Indeed, we found that WNT/FGF-induced TBXT^low^ cultures consisted predominantly of SOX2^+^ (∼90% of total) cells, most of which expressed CDX2 and HOXC9 (**Fig. 2A, 2B**). Transcriptome profiling following RNA sequencing confirmed that pro-mesodermal/NMP markers (*TBXT, TBX6, MIXL1, EVX1, MSGN1*) were significantly downregulated (≥2 fold, FDR ≤ 0.05) or extinct in D7 cultures compared to D3 NMP-like cells (**Fig. 2C, Table S1).** However, the expression of NMP-associated genes linked to a subsequent pre-neural/early spinal cord identity (*NKX1-2, CDX1/2*) remained at high levels (**Fig. 2C)** and transcripts associated with neural lineage specification (e.g. *NEUROG2, ASXL3, POU3F2, ASCL1, IRX3, PAX6)* (Bosse et al., 1997; Delfino-Machin et al., 2005; Delile et al., 2019; Ericson et al., 1997; Lichtig et al., 2020; Nordstrom et al., 2006; Sagner et al., 2018; Sommer et al., 1996; Zhao et al., 2014) were significantly upregulated (≥2 fold, FDR ≤ 0.05) relative to D3 NMP-like cells **(Fig. 2D, Table S1**). Moreover, cells were marked by expression of early neural crest/dorsal neural tube/BMP-TGFβ-related transcripts (e.g *NR2F1/2, TGFBI, TGFB2, BMP6, ID4, BMPR1B)* (Frith et al., 2018; Kee and Bronner-Fraser, 2001; Lee et al., 1998; Rada-Iglesias et al., 2012), consistent with a pro-dorsalising role of WNT and FGF signalling pathways and in line with our previous work showing that hPSC-derived NMP-like cultures contain trunk neural crest precursors (Frith et al., 2018) **(Fig. 2D, Table S1**). Addition of exogenous RA, which has been shown to promote neural differentiation of NMPs and their pre-neural derivatives, both *in vivo* and *in vitro* (Diez del Corral et al., 2002; Diez del Corral et al., 2003; Gouti et al., 2017; Gouti et al., 2014; Verrier et al., 2018), in the presence of FGF2 and CHIR, boosted further the expression of the definitive neural marker *PAX6* and diminished the levels of the NMP/pre-neural marker *NKX1-2* without affecting *CDX2* expression (**Fig. S2**). Finally, although induction of *HOX* transcripts (especially those belonging to the *HOXB* cluster) was initiated in D3 NMP-like cells, their levels continued to be upregulated so that by day 7 of differentiation most members of PG (1-9) were highly expressed (**Fig. 2E**). Collectively, these data indicate that a combination of WNT and high FGF signalling activities steer NM-potent spinal cord progenitors toward a pre-neural/early neural state exhibiting dorsal neural tube features while simultaneously promoting a gradually more posterior axial identity.

**Figure 2.**
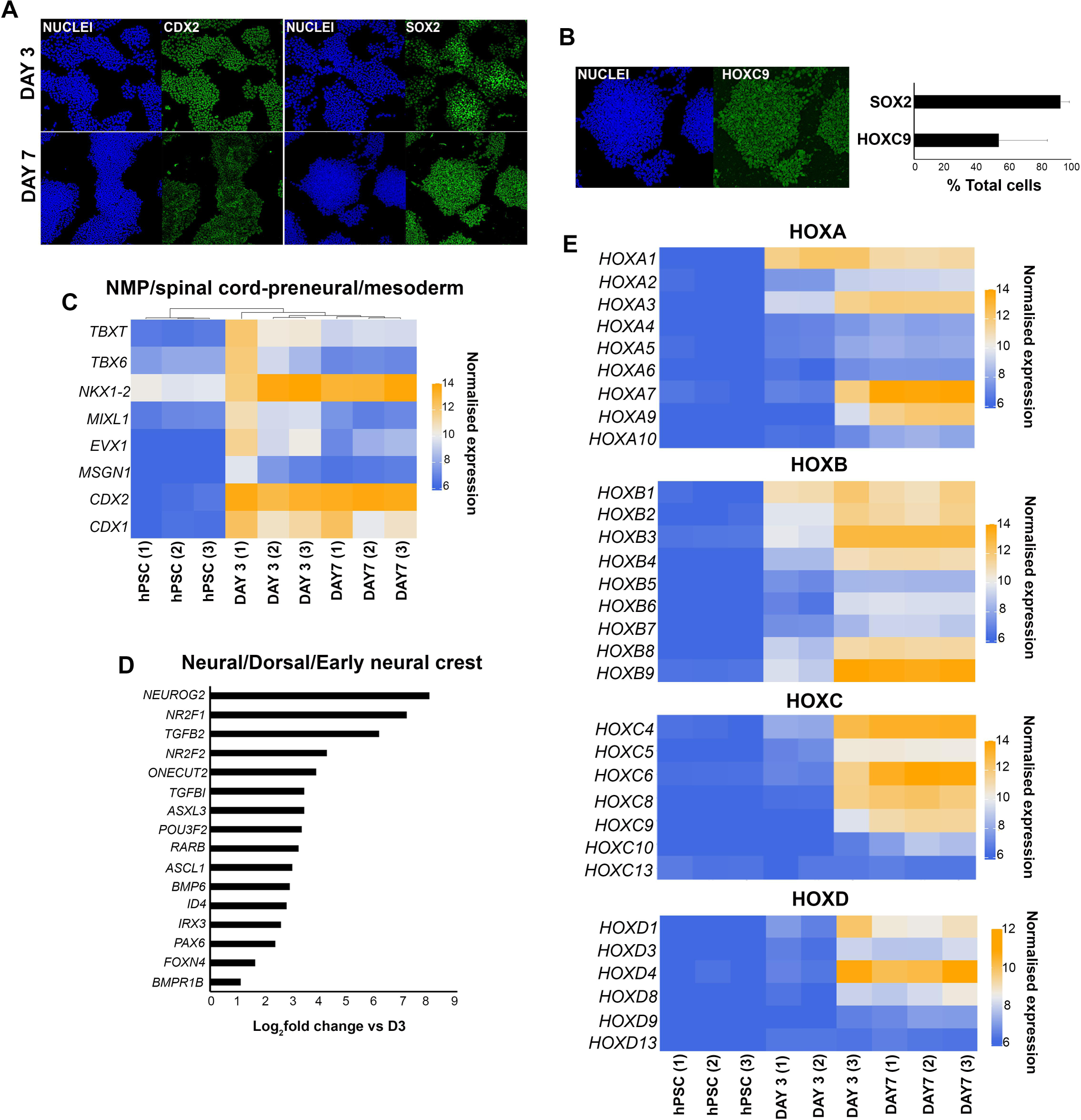
WNT-FGF-mediated induction of posterior neurectoderm from human NMP-like cells. **(A)** Immunofluorescence analysis of the expression of CDX2 and SOX2 at D3 (=NMP-like cells) and D7 early neurectoderm cells following differentiation of hPSCs in the presence of 3 µM CHIR and 100 ng/ml FGF2 as described in Fig. 1A. **(B)** Immunofluorescence analysis of the expression of HOXC9 expression in D7 WNT-FGF-induced early neurectoderm cells following differentiation of hPSCs. The quantification of SOX2^+^ and HOXC9^+^ cells following immunostaining and image analysis is shown below. Average percentages of cells over total numbers are depicted. The data in the graph were obtained after scoring 5-20 random fields per experiment (three independent replicates) i.e. a total of 42 fields for three experiments and the error bars/standard deviation represent the variation across all fields and experiments. Error bars=S.D. **(C)** Heatmap showing the normalised expression values of NMP/early spinal cord/paraxial mesoderm transcripts in three independent hPSC, NMP-like (day 3) and WNT-FGF-induced early neurectoderm (day 7) sample replicates. Numbers in brackets indicate individual replicates. Values were variance stabilised using the vst function in DESeq2. **(D)** Log fold induction of representative neural/dorsal/BMP/TGFβ-associated markers in D7 WNT-FGF treated cultures compared to D3 NMP-like cells. **(E)** Heatmaps showing the normalised expression values of *HOX* transcripts belonging to different clusters in three independent hPSC, NMP-like (day 3) and WNT-FGF-induced early neurectoderm (day 7) sample replicates. Numbers in brackets indicate individual replicates. Values were variance stabilised using the vst function in DESeq2.

### Efficient induction of a ventral identity in NMP-derived neurectoderm requires combined BMP-TGFβ signalling pathway inhibition

We next tested culture conditions for the efficient ventralisation of early dorsal posterior neurectoderm derived from human NMP-like cells following WNT and FGF stimulation. *In vivo*, the ventral neural tube, where MNs and their progenitors arise, is patterned by a gradient of notochord/floor plate-derived SHH signalling activity (reviewed in (Sagner and Briscoe, 2019). Therefore, we initially treated hPSC-derived NMP-like cells with WNT-FGF agonists and RA to induce early posterior neurectoderm, as described above (**Fig. 2, S2**), together with the SHH signalling pathway agonists SAG and purmorphamine to promote ventralisation (Amoroso et al., 2013) (**Fig. 3A**). We also included a BMP pathway inhibitor (LDN 193189) during the initial induction of NMP-like cells from hPSCs (D0-3) to suppress emergence of neural crest cells as we previously showed that endogenous BMP activity during that stage can promote early neural crest features at the expense of central nervous system neural cells (Frith et al., 2018). Following WNT-FGF agonist removal after D10, cells were further cultured in neurotrophic media containing BDNF, GDNF, L-ascorbic acid and the NOTCH/ γ-secretase inhibitor DAPT (after D14), to promote exit from a progenitor state and MN maturation as described previously (Maury et al., 2015) (**Fig. 3A**). This approach (tested in both human induced PSC (hiPS) and human embryonic stem cell (hES) lines) gave rise to cells defined by the expression of ventral spinal cord/MN progenitor regulators OLIG2, NKX6.1 and NKX6.2 (Briscoe et al., 2000; Lu et al., 2002; Novitch et al., 2001; Sagner et al., 2018; Sander et al., 2000; Takebayashi et al., 2002; Vallstedt et al., 2001) from around D14, as well as the definitive MN markers ISL1/2 (Ericson et al., 1992; Thaler et al., 2004) and MNX1 (Arber et al., 1999; Thaler et al., 1999) around D24, both at the transcript (black bars in graphs in **Fig. 3B)**, and protein level (**Fig. 3C**; black bars in **Fig. 3D**). Differentiating cells retained high levels of HOXC9 expression (∼80% of total) (**Fig. 3D**) indicating the stable acquisition of a posterior axial identity. Together these results suggest that WNT-FGF-induced NMP-derived posterior neurectoderm can be ventralised and give rise to neurons corresponding to those found at thoracic levels of the A-P axis. However, both D14 and D24 cultures were found to be heterogeneous containing only low numbers of OLIG2^+^/ISL1/2^+^/MNX1^+^ MN progenitors/MNs (≤5% of total) (**Fig. 3C, 3D**).

**Figure 3.**
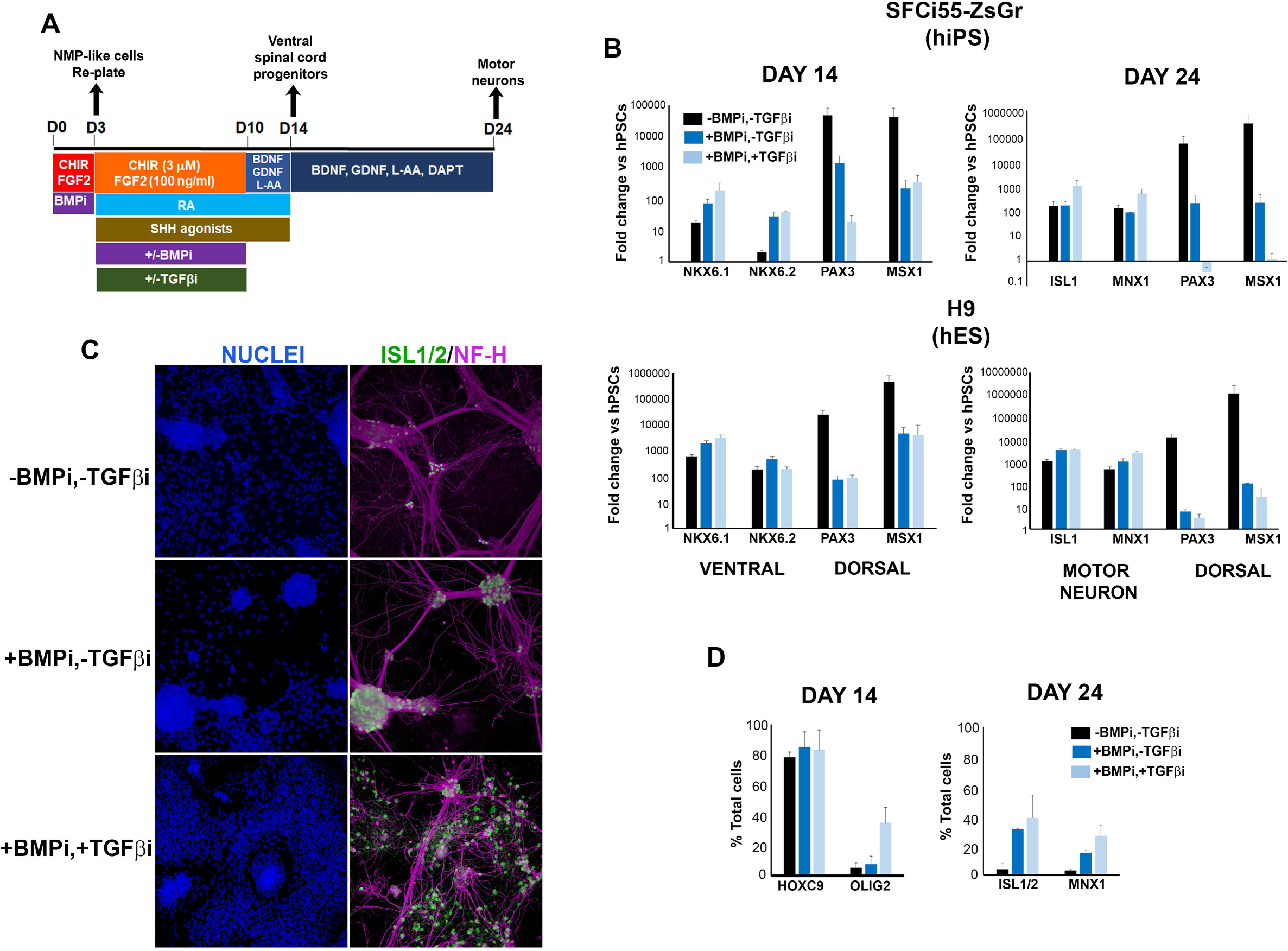
Combined TGFβ/BMP signalling inhibition drives efficient induction of a ventral identity in NMP-derived neurectoderm. **(A)** Diagram depicting the culture conditions employed and tested for the generation of motor neuron (MN) progenitors (D14) and MNs (D24) from hPSC-derived NMP-like cells. **(B)** qPCR expression analysis of indicated dorsal, ventral and MN markers in the absence (black bars) and presence of BMP only (dark blue bars) or TGFβ and BMP (light blue bars) inhibitors (i) at different time points during the differentiation of NMP-like cells to MNs as described in A. Results are shown for a hiPSC (top) and a hESC (bottom) line. Error bars=S.D. (n=3). **(C)** Immunofluorescence analysis of the expression of the MN marker ISL1/2 together with the pan-neuronal marker Neurofilament Heavy Chain (NF-H) in D24 cultures following differentiation of hPSCs in the absence and presence of BMP only or TGFβ and BMP inhibitors. **(D)** Quantification of cells marked by expression of indicated proteins at D14 and D24 after MN differentiation of hPSCs in the absence (black bars) and presence of BMP only (dark blue bars) or TGFβ and BMP (light blue bars) inhibitors and following immunofluorescence and image analysis. The data in the graph were obtained after scoring: D14: two random fields per experiment (two independent replicates) (for - TGFβi,-BMPi and -TGFβi,+BMPi) or four random fields per experiment (two independent replicates) (for +TGFβi, +BMPi); D24 three random fields per experiment (two independent replicates) (for -TGFβi,-BMPi and -TGFβi,+BMPi) or five random fields per experiment (two independent replicates) (for +TGFβi, +BMPi). Error bars = s.d.

The low efficiency of our MN differentiation protocol prompted us to analyse in detail the composition of the “contaminating” non-MN cell types present in the cultures. Given the reported propensity of NMP-like cells to generate neural crest (Frith et al., 2018; Hackland et al., 2019) and the dorsal character of NMP-derived early neurectoderm (**Fig. 2D**), we tested the presence of dorsal neural cell types during MN differentiation. Indeed, qPCR analysis of D14 and D24 hiPS- and hES cell-derived cultures showed that these exhibited very high expression of the dorsal neural tube /neural plate border-early neural crest markers *PAX3* and *MSX1* (Goulding et al., 1991; Timmer et al., 2002) (**Fig. 3B**, black bars). This finding indicates that stimulation of SHH signalling alone was not sufficient to eliminate the dorsal bias of human NMP neural derivatives and drive homogeneous ventralisation/induction of MNs.

We hypothesised that active repression of dorsalisation may be required in order to block the induction of unwanted dorsal neural/neural crest cells. To this end, we tested the effect of inhibitors of established dorsalising signalling pathways such as BMP and TGFβ (**Fig. 3A**), components of which were also found to be enriched in D7 FGF-WNT-induced neurectoderm (**Fig. 2D**). Although BMP antagonism alone was found to be effective (dark blue vs black bars in **Fig. 3B; Fig. 3C**), inclusion of inhibitors of both pathways, during early neural differentiation of hPSC-derived NMP-like cells (D3-10), was the optimal approach for eliminating the expression of *PAX3* and *MSX1* by D24 (light blue bars in **Fig. 3B**) and improving the induction of ventral/MN progenitor markers *NKX6.1*/*NKX6.2* and *ISL1/MNX1* at D14 and 24, respectively (light blue bars in **Fig. 3B**). A similar positive effect of dual BMP-TGFβ inhibition on the overall numbers of OLIG2 and ISL1/2/MNX1 protein-expressing cells was also observed (6-10-fold increase compared to untreated controls while remaining positive for HOXC9) (**Fig. 3C**; light blue bars in **Fig. 3D**). We conclude that the simultaneous blocking of BMP/TGFβ signalling activities, combined with stimulation of SHH signalling, drive efficient acquisition of a ventral identity by NMP-derived posterior spinal cord progenitors *in vitro*.

### Generation of functional posterior MNs from hPSC-derived NMP-like cells

Using the above findings, we established an improved protocol for the *in vitro* generation of posterior MNs. This relies on the induction of early posterior neurectoderm (via WNT-FGF-RA stimulation), which is simultaneously ventralised at high efficiency (SHH stimulation/BMP-TGFβ inhibition), from NMP-like cells, followed by culture in neurotrophic media (BDNF, GDNF, L-ascorbic acid, DAPT) as described above (**Fig. 3A**). We tested this protocol in four different hPSC lines (two hiPSCs: SFCi55-ZsGr, MIFF1 and two hESCs: H7 and H9 (Desmarais et al., 2016; Lopez-Yrigoyen et al., 2018; Thomson et al., 1998)). We found that in all lines tested, early ventral spinal cord/MN progenitor-related transcripts (*NKX6-1, OLIG2*) were induced between D7-14 and, concurrently, a wave of expression of committed MN markers (*MNX1, ISL1*) occurred primarily during the D14-24 time window (**Fig. 4A**). The transcript levels of the early spinal cord progenitor marker *CDX2* became extinct by D24 and posterior HOX transcripts belonging to PG(6-10) were highly expressed throughout differentiation (**Fig. 4A**). It should be noted that the temporal expression dynamics and extent of induction of some of these genes varied between different hPSC lines (**Fig. 4A**), possibly reflecting the well-described transcriptional and epigenetic differences of hPSC lines that affect their differentiation potential (Bock et al., 2011; International Stem Cell, 2018; Koyanagi-Aoi et al., 2013; Nishizawa et al., 2016). Collectively, these results indicate that our protocol promotes the progressive and robust induction of ventral spinal cord progenitors and MNs exhibiting a posterior axial identity.

**Figure 4.**
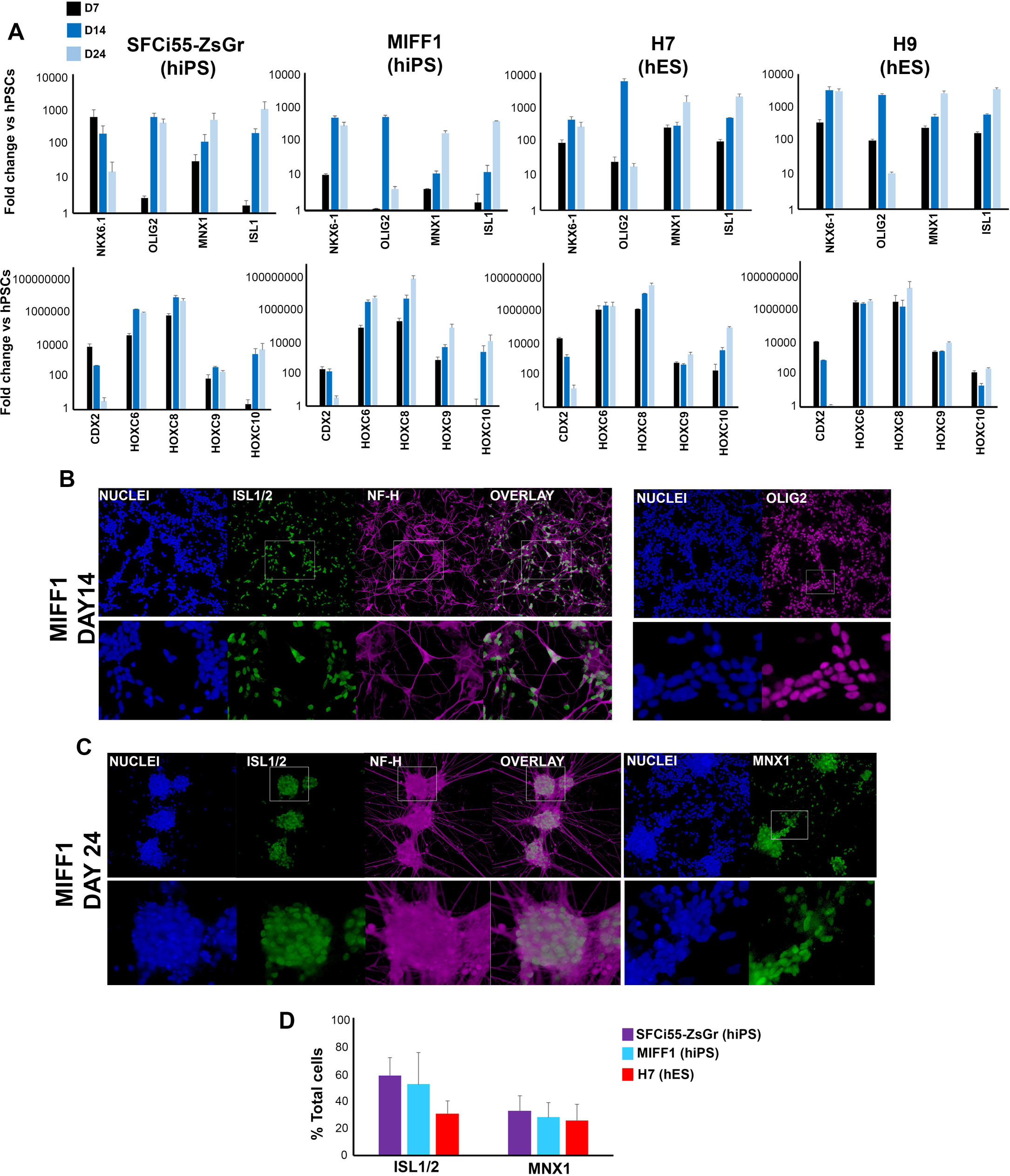
Generation of posterior motor neuron progenitors and motor neurons from hPSCs. **(A)** qPCR expression analysis of indicated MN progenitor/MN markers (top) and posterior *HOX* transcripts (bottom) at different time points during the differentiation of various human induced pluripotent (hiPS) and embryonic stem cell (hES) lines to MNs in the presence of BMP/TGFβ inhibitors as described in Fig. 3A. Error bars=S.D. (n=3). **(B)** Immunofluorescence analysis of the expression of the MN marker ISL1/2 together with the pan-neuronal marker Neurofilament Heavy Chain (NF-H), and the MN progenitor marker OLIG2 in day 14 cultures differentiated from MIFF1 hiPSCs in the presence of BMP/TGFβ inhibitors as described in Fig. 3A. Magnified regions corresponding to the insets are also shown. **(C)** Immunofluorescence analysis of the expression of the MN marker ISL1/2 and NF-H in day 24 cultures differentiated from MIFF1 hIPS cells in the presence of BMP/TGFβ inhibitors as described in Fig. 3A. Magnified regions corresponding to the insets are also shown. **(D)** Quantification of cells marked by expression of MN-associated proteins at D24 after differentiation of indicated hPSC lines in the presence of BMP/TGFβ inhibitors as described in Fig. 3A, and following immunofluorescence and image analysis. The data in the graph were obtained after scoring seven random fields per experiment (three independent replicates). Error bars = s.d.

To further verify the identity of the differentiating cells we carried out antibody staining and fluorescence microscopy. These analyses confirmed that D14 cultures were predominantly HOXC9^+^ (∼80% of total cells) and consisted of a mix of OLIG2^+^ MN progenitors (35-70% of total cells depending on the hPSC line) and ISL1/2-Neurofilament Heavy Chain (NF-H)-double positive MNs (∼30-40% of total cells depending on the hPSC line) (**Fig. 3D, 4B, S3A, S3C** and data not shown). At D24 we observed the emergence of a considerable number of 3-dimensional clusters consisting of ISL1/2^+^ cells (∼30-60% of total cells depending on the hPSC line) and/or MNX1^+^ cells (25-35% of total cells) forming networks via NF-H^+^ neuronal connections (**Fig. 3C, 3D, 4C, 4D, S3B**). The lower number of MNX1^+^ cells in our cultures, compared to those obtained with protocols generating MNs of a more anterior axial identity (Du et al., 2015; Maury et al., 2015), is likely to reflect the presence of thoracic MNs exhibiting a preganglionic column (PGC) character, which is marked by the absence of MNX1 expression (William et al., 2003). Moreover, we found that D14 MN progenitor/MN cutlures retained the expression of all appropriate markers and could be differentiated into more mature MNs following freezing and thawing (**Fig. S3D**). Taken together, these data suggest that our protocol can be applied to various hPSC lines to generate considerable numbers of posterior MN progenitors/MNs via an NMP-like intermediate.

We next tested the ability of posterior D14 MN progenitors/early MNs to integrate within the embryonic spinal cord and establish neuronal projections toward muscles *in vivo.* Such a population could be a promising platform for the development of cell therapy-based strategies against ALS and spinal cord injury (Kadoya et al., 2016; Kondo et al., 2014; Kumamaru et al., 2018; Zalfa et al., 2019). We employed cells derived from SFCi55-ZsGr hiPSCs, which carry a constitutive ZsGreen fluorescent reporter (Lopez-Yrigoyen et al., 2018), and transplanted them into chick embryos, a well-established method for testing the developmental potential of PSC-derived cells in an embryonic environment (Frith et al., 2018; Peljto et al., 2010; Wichterle et al., 2002) (**Fig. S4A**). Donor cells were grafted to the posterior open neural tube of Hamburger and Hamilton (HH) stage 10-13 chick embryos i.e. a site that gives rise to brachial/thoracic regions of the spinal cord (**Fig. 5A**). This host site can therefore be defined as “homotopic” in terms of axial identity, for the posterior MN progenitors/MN donor cells. After incubation, host embryos were harvested at HH26-30. 22/30 embryos showed no developmental abnormalities and 11 of these contained ZsGREEN^+^ cells located within the thoracic spinal cord between the forelimb and hindlimb levels, as revealed by wholemount imaging (**Fig. 5A**). Embryos were sectioned (n=8) and the incorporation/differentiation of hPSC-derived fluorescent cells was assessed following immunostaining and fluorescence microscopy.

**Figure 5.**
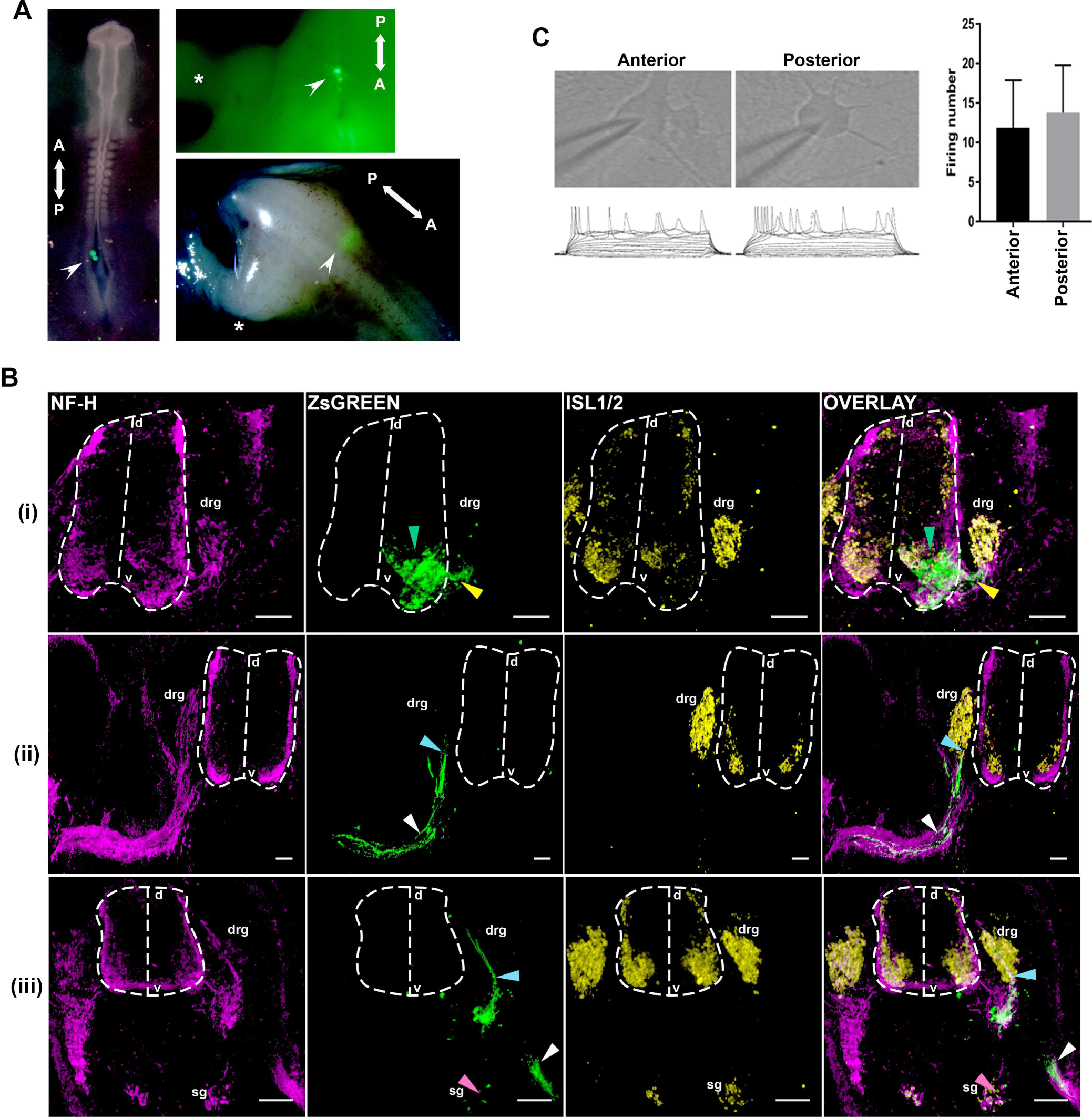
Testing the functionality of hPSC-derived posterior MNs. **(A)** Whole mount embryo images showing the engrafted ZsGREEN+ donor cells (indicated by arrow head) straight after transplantation (left) and prior to sectioning, following host embryo incubation (right). Asterisks indicate the position of the limbs. A, anterior; P, posterior. **(B)** Representative immunofluorescence analysis images of ZsGREEN, NF-H and ISL1/2 expression marking donor human cells, neuronal projections and dorsal root ganglion (drg)/MN-containing ventral neural tube/sympathetic ganglia (sg) respectively, in sections of HH26-30 chick embryos grafted with hPSC-derived ZsGREEN+ posterior D14 MN progenitor/MN cultures. Dotted outlines mark the boundaries of the neural tube. D, dorsal; v, ventral. Green arrowheads mark cell integration within the ventral MN progenitor domain. Yellow arrowheads mark ZsGREEN-positive axons exiting the neural tube via the MN ventral root. White arrowheads mark ZsGREEN-positive axons exiting the neural tube and projecting along the MN ventral root towards muscle target cells, following the endogenous pathway. Pink arrowheads mark ZsGREEN+ cells within the sympathetic chain ganglia. Blue arrowheads mark donor cell projections toward the dorsal root ganglia. **(C)** Firing rate of anterior and posterior hPSC-derived MNs. Whole-cell current clamp recordings were performed at room temperature and injected with 15 steps of currents from -80pA to +60pA (10pA increase for each step). Representative traces of each group are shown on the left. Total firing number of each neuron with magnitude of 20mV or higher were counted for analysis. Analysis results were shown on the right panel. (No. of recorded neurons =14, mean ± SEM; one-way ANOVA Tukey’s multiple comparison test, no significant difference between the two groups).

We found that the human donor cells (6/8 embryos) integrated well within the ventral part of the neural tube encompassing the MN progenitor domain (**Fig. 5Bi;** green arrowhead) and expressed ISL1/2, HOXC9 and NF-H (**Fig. S4B, C**) consistent with thoracic MN differentiation, although OLIG2+ donor cells were also detected within donor cell clumps (**Fig. S4B**) indicating the presence of more immature MN progenitors as well. Importantly, donor ZsGREEN+ cells projected axons in a manner similar to endogenous host MNs:

i. In 6/8 embryos, ZsGREEN-positive axons exited the neural tube via the MN ventral root (e.g. **Fig. 5Bi;** yellow arrowhead).
ii. In many cases (4/8 embryos), sections adjacent to those with the main bulk of integrated cells revealed donor cell-derived axons exiting the neural tube and projecting along the MN ventral root towards muscle target cells, following the endogenous pathway (**Fig.5Bii, Biii**; white arrowheads).
iii. In 2/8 embryos we found ZsGREEN+ axons exiting the ventral neural tube and projecting toward the sympathetic chain ganglia, a feature of PGC MNs emerging at the thoracic levels of the spinal cord (**Fig. 5Biii;** pink arrowhead).

Interestingly, we also observed donor cell projections toward (and sometimes into/beyond) the dorsal root ganglia (drg) following exit via the ventral root (5/8 embryos; **Fig. 5Bii, Biii;** blue arrowheads). Collectively, these findings suggest that thoracic MN progenitors/MNs generated from hPSC-derived NMP-like can integrate efficiently into the posterior embryonic spinal cord following homotopic transplantation to the chick embryo, and establish appropriate neuronal projections.

We next examined the influence of axial identity on the electrophysiological properties of spinal cord MNs by comparing our posterior MN cultures with their anterior counterparts, since the firing rate of various MN subtypes has been linked to differential vulnerability to ALS (Brockington et al., 2013; Nijssen et al., 2017). To this end, we generated MNs of a hindbrain/cervical character using an established differentiation strategy: hPSCs were first treated with TGFβ/BMP signalling inhibitors to induce an anterior *Hox*-negative neurectodermal intermediate, and then RA/SHH agonists to promote posteriorisation/ventralisation (Amoroso et al., 2013; Chambers et al., 2009; Wichterle et al., 2002) (**Fig. S5A**). Following culture in neurotrophic media, the resulting cultures exhibited mixed expression of MN progenitor (*NKX6.1*/*OLIG2*) and committed MN (*ISL1*) markers at D14 of differentiation (**Fig. S5B**) while at D24 they consisted mainly of ISL1/2^+^ clusters with NF-H^+^ neuronal projections (**Fig. S5C**).The cells were also found to exhibit high levels of *HOX* PG (1-5) gene expression and low levels of *MNX1* as well as no/very low *CDX2* and *HOX* PG (6-10) gene expression, confirming their hindbrain/cervical axial identity (**Fig. S5B**). Whole-cell current clamp recordings revealed that both these anterior MNs and their posterior thoracic counterparts, generated using our protocol, exhibited comparable action potential firing rates (**Fig. 5C)**. These were similar to those reported for *in vivo-*transplanted mouse PSC-derived MNs (Soundararajan et al., 2006) and rat embryonic MNs (Alessandri-Haber et al., 1999), suggesting that both anterior and posterior hPSC-derived MNs are equivalent in terms of electrophysiology.

## Discussion

Here, we have described an optimised protocol for the generation of functional posterior MNs from various hPSCs via the induction of NMP-like cells. Our work addresses a major gap in conventional hPSC differentiation strategies, namely the low efficiency of most protocols in generating thoracic spinal cord cells and MNs. It therefore opens new avenues toward the development of novel regenerative medicine and disease modelling approaches that rely on the induction of defined regional identities *in vitro*. Our approach gives rise to considerable numbers of OLIG2^+^ MN progenitors and ISL1/2^+^ MNs (up to 70% and 60% of total cells respectively), which also express high levels of HOX PG (6-9) genes thus reflecting their posterior brachial/thoracic axial identity. These yields represent an improvement over previous protocols for the production of thoracic MNs from adherent human NMP-like cells (Estevez-Silva et al., 2018; Lippmann et al., 2015) and this is, to our knowledge, the first time that the NMP-based, *in vitro* generation of posterior human MNs is characterised in detail and across different hPSC lines.

We found that the posterior MNs produced using our approach are functional in terms of electrophysiology, behaving similarly to hPSC-derived MNs exhibiting a hindbrain/anterior spinal cord character. Moreover, following transplantation to the posterior spinal cord of chick embryos, our posterior MNs showed hallmarks of their *endogenous in vivo* counterparts i.e. expression of ISL1/2 and HOXC9 and establishment of NF-H-positive axonal projections exiting, via the ventral root, toward target muscles/sympathetic ganglia. Interestingly, grafted donor cells also tended to form neuronal projections toward the DRG, a pattern that has been previously observed with grafts of PSC-derived MNs of a more anterior (hindbrain/cervical/anterior brachial) axial identity (Amoroso et al., 2013; Soundararajan et al., 2006) indicating that such cells may also be present in our cultures. Alternatively, this may reflect low levels of CXCR4 expression in our donor D14 posterior MN progenitor/MN cultures due to incomplete differentiation, as similar projections of ventral spinal cord MNs toward the DRG have been previously reported in *Cxcr4* mutant mice (Lieberam et al., 2005).

Our data show that distinct levels of WNT and FGF signalling activities steer hPSC-derived NMP-like cells toward their immediate derivatives paraxial mesoderm or early neurectoderm. The transition to the latter entity occurs via a pre-neural NKX1-2^+^SOX2^+^CDX2^+^ state and day 7 WNT-FGF-induced cultures are composed by a mixture of such pre-neural and early neural progenitors. Their emergence is likely to be driven directly by FGF and WNT signalling, as these pathways have been shown to promote specification and maintenance of early posterior progenitors in the caudal lateral epiblast/pre-neural tube *in vivo* (Diez del Corral et al., 2002; Diez del Corral et al., 2003; Mathis et al., 2001; Nordstrom et al., 2006; Olivera-Martinez et al., 2014; Olivera-Martinez and Storey, 2007; Storey et al., 1998; Takemoto et al., 2006).

Previous findings have suggested that spinal cord neurectoderm can be induced by addition of RA to WNT-FGF-induced hPSC-derived NMP-like cells: the longer the duration of WNT-FGF treatment during NMP induction, the more posterior the axial identity of their RA-induced neural derivatives (Lippmann et al., 2015). Similarly, we show that sequential *HOX* gene activation and hence the acquisition of a posterior axial identity is a function of the duration of WNT-FGF stimulation. However, we also found that progressive *HOX* gene induction takes place during the transition of NMP-like cells toward a pre-neural/early neural state rather than within a homogeneous TBXT^+^SOX2^+^ population maintained by continuous WNT-FGF treatment. Collectively, these findings suggest that WNT and FGF stimulation can “program” a posterior axial identity (marked by posterior *HOX* gene expression) independently of the lineage identity of the cells these pathways act on.

The simultaneous induction of both a posterior axial and early neural identity, following WNT-FGF treatment of human NMP-like cells, is also marked by upregulation of dorsal neural tube markers and high activity of dorsalising BMP/TGFβ signalling pathways, in agreement with previous studies examining neural differentiation of hPSC-derived NMP-like cells (Denham et al., 2015; Frith et al., 2018; Hackland et al., 2019; Verrier et al., 2018). Both WNT and FGF pathways have been implicated in the promotion of a dorsal identity/repression of ventral patterning genes in neural cells (Alvarez-Medina et al., 2008; Diez del Corral et al., 2003; Muroyama et al., 2002) and our data suggest that their pro-dorsal effect may be mediated by activating downstream BMP and TGFβ signalling cascades. Furthermore, an increase in the expression of BMP and TGFβ pathway components has been reported to coincide with the onset of neuronal differentiation in the posterior neural tube during chick embryonic development (Olivera-Martinez et al., 2014). Hence, neuralisation and acquisition of a dorsal identity may be tightly coupled events during the generation of posterior neurectoderm *in vitro* and presage ventralisation. The latter can only be achieved by combining SHH agonist treatment with dual BMP-TGFβ signalling inhibition reflecting *in vivo* data showing that MN specification also depends on the activity of BMP-TGFβ signalling antagonists such as Noggin and Follistatin (Liem et al., 2000; McMahon et al., 1998).

To date, it has been unclear whether the axial identity of neural derivatives of PSCs shapes their *in vivo* functionality, a critical parameter to consider in the development of cell replacement therapies. Previous studies employing transplantation of PSC-derived neural progenitors of a presumed anterior axial identity, have suggested that donor cells can survive, differentiate and even promote functional recovery in rodent disease and injury models within posterior spinal cord locations (Iwai et al., 2015; Kondo et al., 2014; Lu et al., 2017; Lu et al., 2014; Tsuji et al., 2010; Yohn et al., 2008). However, more recent data based on the use of both *in vivo* and *in vitro*-derived neural progenitors have demonstrated that axial level homology between donor cell and spinal cord transplantation sites is a key factor for functional reconstitution in the injured spinal cord (Kadoya et al., 2016; Kumamaru et al., 2018). We envisage that the efficient generation of posterior spinal cord cells using our protocol will facilitate their direct functional comparison to their anterior counterparts, thus revealing whether the *in vivo* behaviour of hPSC-derived derived spinal cord cells is intrinsically determined by their *in vitro* axial identity, or plastic and influenced by extrinsic factors.

## Methods and Materials

### Cell culture and differentiation

Use of hES cells has been approved by the Human Embryonic Stem Cell UK Steering Committee (SCSC15-23). The following hPSC lines were employed: SFCi55-ZsGr (Lopez-Yrigoyen et al., 2018), MIFF1 (Desmarais et al., 2016), H7 and H9 (Thomson et al., 1998). All cell lines were tested mycoplasma negative. Cells were cultured in feeder-free conditions in Essential 8 (Thermo Fisher or made in-house) medium on laminin 521 (Biolamina). For NMP-like cell differentiation, hPSCs (60-80% confluency) were dissociated using PBS/EDTA and plated at a density of 63,000 cells/cm^2^ on vitronectin (Thermo Fisher)-coated wells, directly into N2B27 medium containing CHIR99021 (3 µM, Tocris), FGF2 (20 ng/ml, R&D), LDN-193189 (100 nM, Tocris), and ROCK inhibitor Y-27632 (10 µM, Adooq Biosciences) for the first only day. For early neurectoderm differentiation, day 3 hPSC-derived NMP-like cells were dissociated using Accutase and re-plated at a density of 37,500-86,000 cells /cm^2^ on Geltrex (Thermo Fisher)-coated plates directly in N2B27 containing CHIR99021 (3 µM, Tocris) and FGF2 (100 ng/ml, R&D) in the presence or absence of retinoic acid (0.1 µM, Tocris). For paraxial mesoderm differentiation, day 3 NMP-like cultures were treated with Accutase and replated at a density of 45,000 cells/cm^2^ on Geltrex-coated plates in N2B27 containing FGF2 (40 ng/ml, R&D) and CHIR99021 (8 µM, Tocris) for two days. For posterior MN differentiation, day 3 NMP-like cell cultures were treated with Accutase and replated at a density of 63,000 cells//cm^2^ in N2B27 supplemented with CHIR99021 (3 µM, Tocris), FGF2 (100 ng/ml, R&D), Purmorphamine (1µM, Sigma), SAG (0.5 µM, Tocris), all-trans retinoic acid (0.1 µM, Tocris), LDN-193189 (100 nM, Tocris), DMH1 (1µM, Tocris), SB431542 (10µM, Tocris) and ROCK inhibitor Y-27632 (10 µM, Adooq Biosciences) for the first only day, as indicated, on Geltrex-coated plates. Media were replaced every 2 days. Cells were re-plated on day 7 of differentiation using Accutase to lift and dissociate and seeded at 86,000 cells/cm^2^, on Geltrex-coated plates, in the same media as described above. At day 10 of differentiation the media was changed to N2B27 supplemented containing Purmorphamine (1µM, Sigma), SAG (0.5 µM, Tocris), all trans retinoic acid (0.1 µM, Tocris), BDNF (20ng/ml, Peprotech), GDNF (20ng/ml, Peprotech) and L-Ascorbic Acid (200 µM, Sigma) and replaced again at day 12. At day 14 differentiation, cells were re-plated using Accutase and seeded at a density of 100,000 cells/cm^2^ onto Geltrex-coated plates in N2B27 medium supplemented with BDNF (20ng/ml, Peprotech), GDNF (20ng/ml, Peprotech) and L-Ascorbic Acid (200 µM, Sigma), DAPT (10µM, Sigma) ROCK inhibitor Y-27632 (10 µM, Adooq Biosciences) for the first only day. Media were replaced every other day until day 24 or 36 of differentiation. For anterior MN differentiation, hPSCs (confluency of 60-80%) were seeded at 63,000 cells/cm^2^, on Geltrex-coated plates, in N2B27 containing LDN-193189 (100 nM, Tocris), SB431542 (10µM, Tocris) and ROCK inhibitor Y-27632 (10 µM, Adooq Biosciences) for the first only day. On day three of differentiation, cells were re-plated at 63,000 cells/cm^2^, on Geltrex coated plates, in N2B27 supplemented with LDN-193189 (100 nM, Tocris), DMH1 (1µM, Tocris), SB431542 (10µM, Tocris), Purmorphamine (1µM, Sigma), SAG (0.5 µM, Tocris), all trans retinoic acid (0.1 µM, Tocris), and ROCK inhibitor Y-27632 (10 µM, Adooq Biosciences) for the first only day. Media were re-placed at day 6 of differentiation. At day 7 cells were re-plated at a density of 86,000 cells/cm^2^ on Geltrex-coated plates, in N2B27 supplemented with LDN-193189 (100 nM, Tocris), SB431542 (10µM, Tocris), Purmorphamine (1µM, Sigma), SAG (0.5 µM, Tocris), all trans retinoic acid (0.1 µM, Tocris), and ROCK inhibitor Y-27632 (10 µM, Adooq Biosciences) for the first only day. At day 10 the media was changed to N2B27 supplemented containing Purmorphamine (1µM, Sigma), SAG (0.5 µM, Tocris), all trans retinoic acid (0.1 µM, Tocris), BDNF (20ng/ml, Peprotech), GDNF (20ng/ml, Peprotech) and L-Ascorbic Acid (200 µM, Sigma) and replaced again at day 12. At day 14 differentiation, cells were re-plated using Accutase and seeded at a density of 100,000 cells/cm^2^ onto Geltrex-coated plates in N2B27 medium supplemented with BDNF (20ng/ml, Peprotech), GDNF (20ng/ml, Peprotech) and L-Ascorbic Acid (200 µM, Sigma), DAPT (10µM, Sigma) ROCK inhibitor Y-27632 (10 µM, Adooq Biosciences) for two days. Media were replaced every other day until day 24.

### RNA sequencing

#### Sample preparation

For RNA sequencing, we employed hESCs, D3 NMP-like cells and D7 CHIR-FGF-treated spinal cord progenitors. We used a genetically modified hES cell line of H9 background also containing a Tetracycline inducible shRNA cassette against TBXT (Bertero et al., 2016). Cells were cultured in the absence of Tetracycline and exhibited a differentiation behaviour comparable to unmodified controls (data not shown and (Bertero et al., 2016). Total RNA was harvested using the total RNA purification plus kit (Norgen BioTek) according to the manufacturer’s instructions. Sample quality control, library preparation and sequencing were carried out by Novogene (http://en.novogene.com). Library construction was carried out using the NEB Next® Ultra™ RNA Library Prep Kit and sequencing was performed using the Illumina NoveSeq platform (PE150).

#### Pre-processing of reads

FastQC v0.11.2 (Andrews, 2010) was used for quality control of raw reads. Library adapters were trimmed using Trim Galore v0.4.0 (Krueger, 2012). Paired-end reads were aligned to the human reference genome assembly GRCh38 (Ensembl Build 79) using STAR v2.4.2a (Dobin et al., 2013) in the two-pass mode. Expected gene counts were extracted using RSEM v1.3.0 (Li and Dewey, 2011). Genes expressed in < 3 samples or with total counts ≤ 5 among all samples were removed.

#### Differential expression and visualisation

Differential expression analysis was performed using the DESeq2 (v1.22.2) package (Love et al., 2014) in R (v3.5). Differentially expressed genes with log2FoldChange > |1| and Benjamini-Hochberg adjusted p-value< 0.05 were considered significant. Variance stabilising transformation (vst) (Anders and Huber, 2010) from the DESeq2 package was applied to the read counts for heatmap visualisation. Gene ontology analysis was carried out using the ToppGene suite (https://toppgene.cchmc.org/enrichment.jsp) (Chen et al., 2009).

### Quantitative real time PCR

Total RNA from different samples was harvested using the total RNA purification plus kit (Norgen BioTek) according to the manufacturer’s instructions. First strand cDNA synthesis was performed using the high-Capacity cDNA Reverse Transcription kit (ThermoFisher) and random primers. Quantitative real time PCR was carried out using the Applied Biosystems QuantStudio™ 12K Flex thermocycler together with the Roche UPL system. Statistical significance was calculated using GraphPad Prism (GraphPad Software Inc, USA). Primer sequences are available upon request.

### Immunofluorescence

Cells were fixed for 10 minutes at 4°C in 4% paraformaldehyde (PFA) in phosphate buffer saline (PBS), then washed in PBST (PBS with 0.3% Triton X-100) and treated with 0.5 M glycine/PBST to quench the PFA. Blocking was then carried for 1-3 h in PBST supplemented with 3% donkey serum/1% BSA at room temperature or overnight at 4^0^C. Primary and secondary antibodies were diluted in PBST containing in PBST supplemented with 3% donkey serum/1% BSA. Cells were incubated with primary antibodies overnight at 4°C and with secondary antibodies at room temperature for 2 h in the dark. Cell nuclei were counterstained using Hoechst 33342 (1:1000, Thermo Fisher) and fluorescent images were taken using the InCell Analyser 2500 system (GE Healthcare). Chick embryos were fixed in 4% PFA for 2-3 hours at 4^0^C and left in 30% sucrose solution overnight at 4^0^C. Chick embryos were mounted in OCT (VWR 361603E) and transverse sections (15-20µm) were taken using a cryostat. For immunostaining of sections, overnight incubation with the primary antibody at 4^0^C was followed by short washes in PBS/0.1% Triton X-100 solution (PBST), one hour incubation with the secondary antibody and further PBST washes. Slides were mounted in Fluoroshield with DAPI (Sigma) and imaged on the InCell Analyser 2200 (GE Healthcare) or a Widefield Fluorescent Inverted Microscope (Nikon Eclipse Ti-E). We used the following antibodies: anti-TBXT (1:200; ab209665, Abcam, RRID:AB_2750925), anti-SOX2 (1:200; ab92494, Abcam, RRID:AB_10585428), anti-CDX2 (1:200; ab76541, Abcam, RRID:AB_1523334), anti-ISL1/2 (1:100, DSHB #39.4D5, DSHB, RRID:AB_2314683), anti-anti-Neurofilament, Heavy Chain (1:1000, ab8135, Abcam, RRID:AB_306298), anti-OLIG2 (1:200, AF2418, R&D, AB_2157554), anti-HOXC9 (1:50; ab50839, Abcam, RRID:AB_880494), anti-MNX1 (1:100, DSHB #81.5C10, DSHB, RRID:AB_2145209). Images were processed using Photoshop and Fiji. Nuclear segmentation followed by single cell fluorescence quantification was performed either using CellProfiler (Carpenter et al., 2006) using a custom made pipeline. Cells stained with secondary antibody only were used as a negative control to set a threshold fluorescence intensity. Following nuclear segmentation, the fluorescence intensity of each channel was masked back to nuclei and gave the number of positive (with fluorescence intensity greater than secondary only control) and negative cells per channel.

### Chick embryo grafting

On day 13 of MN differentiation, cells were dissociated and plated at concentrations of 1000-5000 cells/cm^2^ to form hanging drops in 20µl of DMEM F12 media (Sigma) supplemented with N2 supplement (Thermo Fisher), MEM Non Essential Amino Acids (Thermo Fisher), Glutamax (Thermo Fisher). After 24 hours of incubation at 37°C the cells generated clumps with a diameter of ∼50–100 mm. Fertilised Bovan brown chicken eggs (Henry Stewart and Co., Norfolk, UK) were staged according to Hamburger and Hamilton (Hamburger and Hamilton, 1951). Clumps were implanted in the neural tube of HH stage 10-13 chick embryos at the level of the newly developing somites. Following incubation, embryos we harvested at HH26-30.

### Electrophysiology

Neurons were plated onto 13mm coverslips at a density of 50,000 cells per coverslip. All recordings were performed at room temperature and all reagents for solutions were purchased from Sigma. Electrodes for patch clamping were pulled on a Sutter P-97 horizontal puller (Sutter Instrument Company) from borosilicate glass capillaries (World Precision Instruments). Coverslips were placed into a bath on an upright microscope (Olympus) containing the extracellular solution at pH7.4 composing of 150 mM NaCl, 5.4 mM KCl, 2 mM MgCl2, 2 mM CaCl2, 10 mM HEPES, 10 mM Glucose, osmolarity∼305 mOsm/Kg. Whole-cell current clamp recordings were performed using an Axon Multi-Clamp 700B amplifier (Axon Instruments, Sunnyvale, CA, USA) using unpolished borosilicate pipettes placed at the cell soma. Pipettes had a resistance of 4-6MΩ when filled with intracellular solution of 140mM K^+^-gluconate, 10 mM KCl, 1 mM MgCl2, 0.2mM EGTA, 9 mM NaCl, 10 mM HEPES, 0.3 mM Na^+^-GTP, and 3 mM Na^+^-ATP adjusted to 298 mOsm/Kg at pH7.4. For both solutions glucose, EGTA, Na^+^-GTP, and Na^+^-ATP were added fresh on each day of the experiment. To identify neurons present, cells were visualised using the microscope x40 objective, and those with a triangular cell body and processes to indicate neuronal morphology were selected. To measure depolarized evoked action potential firing in the cells using a 15-step protocol for a duration of 500 miliseconds, injecting current from - 80pA, every 10pA. Recordings were acquired at ≥10 kHz using a Digidata 1440A analogue-to-digital board and pClamp10 software (Axon Instruments). Electrophysiological data were analysed using Clampfit10 software (Axon Instruments). Firing magnitude 20mV and higher were included for analysis.

## Acknowledgements

We would like to thank Lesley Forrester (University of Edinburgh) and Konstantinos Anastassiadis (Technische Universität Dresden) for providing the SFCi55-ZsGr and MSGN1-VENUS hPSC lines respectively. We would also like to thank Tom Frith, Bethany James, Sophia Tarelli and Theo Wing for technical assistance. Finally, we are grateful to James Briscoe, Sally Lowell, Celine Souilhol and Filip Wymeersch for critical reading of the manuscript. The authors declare no competing or financial interests.

## Author contributions

**Conceptualisation:** AT, MW; **Investigation:** AT, MW, AG, IM, IG, LB, KN; **Data Curation:** IM, IG; **Formal analysis:** AT, MW, AG, IM, IG, LB, KN; **Supervision:** AT, MP, MRG, IB; **Funding acquisition:** AT, MP, PWA, MG; **Project administration:** AT; **Writing - original draft:** AT, MW; **Writing - review & editing:** AT, MW, AG, IM, IG, LB, IB, KN, MRG, MP, PWA.

## Data availability

Sequencing data have been deposited to the NCBI Gene Expression Omnibus (Accession no: GSE151097)

## Supplementary Figure legends

**Figure S1.**
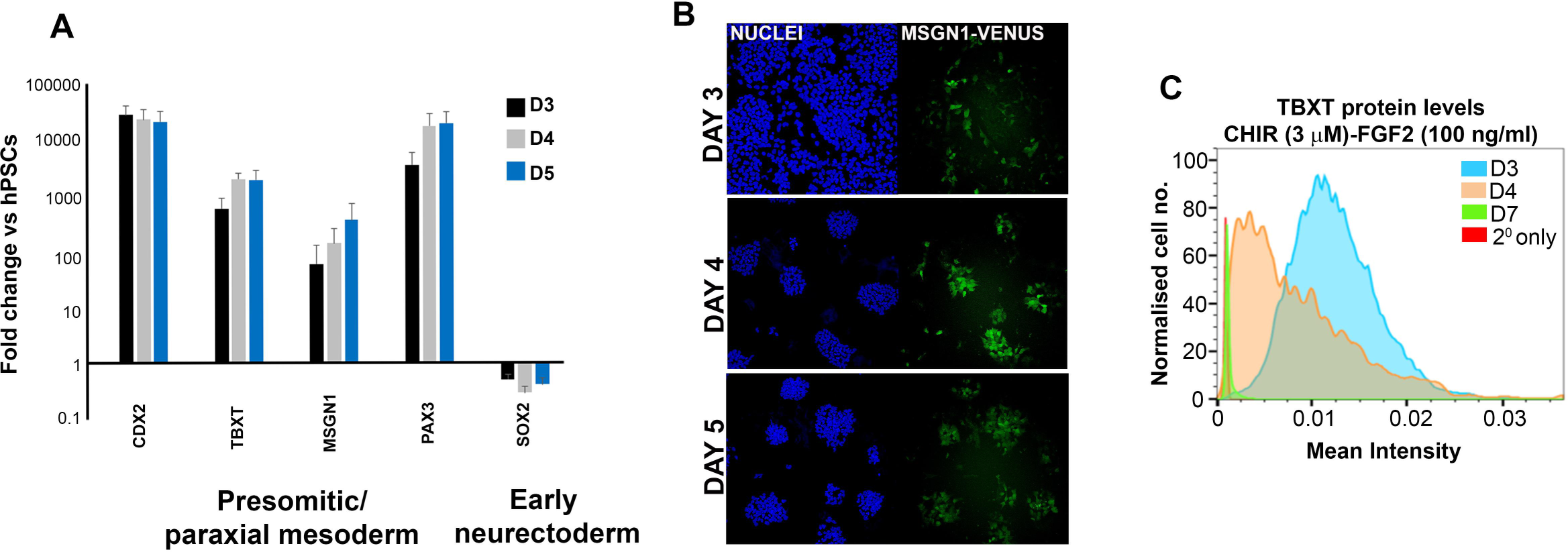
**(A)** qPCR expression analysis of indicated markers at different time points during the differentiation of hPSC-derived NMP-like cells in the presence of 8 µM CHIR and 40 ng/ml FGF2 as described in Fig. 1A. **(B)** Fluorescence analysis of the expression of MSNG1-VENUS at indicated time points during the differentiation of hPSC-derived NMP-like cells (D3) cells in the presence of 8 µM CHIR and 40 ng/ml FGF2 as described in Fig. 1A. Cells were derived from a MSGN1-VENUS reporter hPSC line (Frith et al., 2018). (C) Mean fluorescence intensity of TBXT protein expression in NMP-like (D3) cells and their derivatives during differentiation in the presence of 3µM CHIR and 100 ng/ml FGF2 as described in Fig. 1A.

**Figure S2.**
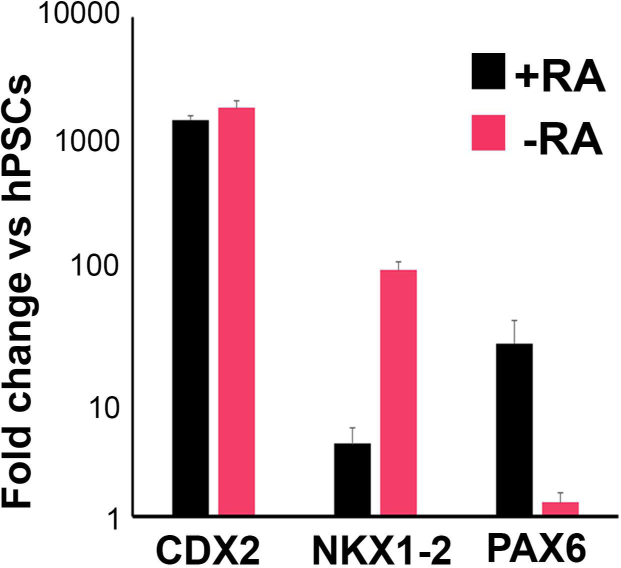
qPCR expression analysis of indicated spinal cord progenitor/neural markers in D7 early neurectoderm cultures induced by WNT and FGF in the presence (+RA) and absence (-RA) of RA.

**Figure S3.**
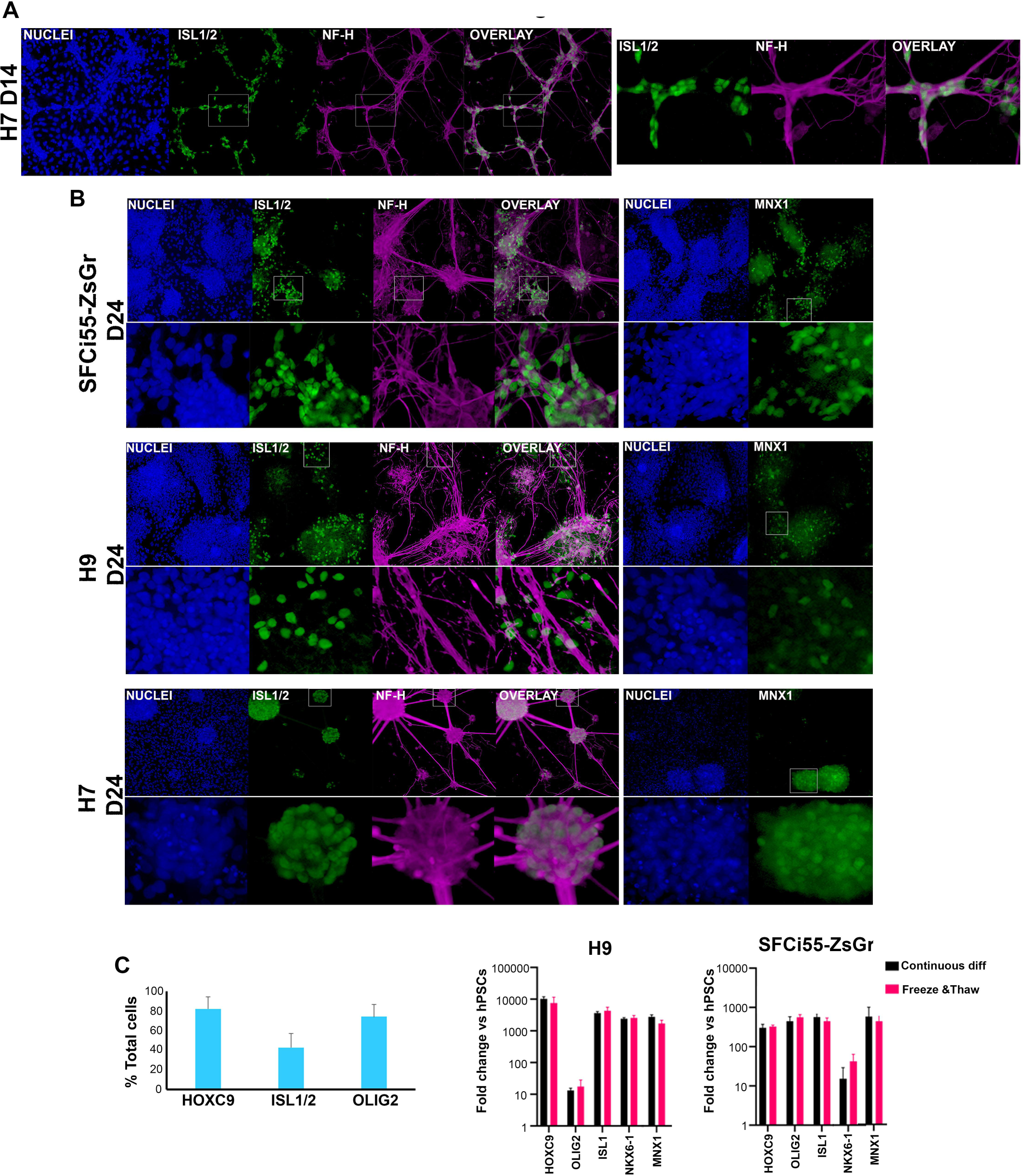
**(A)** Immunofluorescence analysis of the expression of the MN marker ISL1/2 together with the neuronal marker Neurofilament Heavy Chain (NF-H) in day 14 cultures differentiated from H7 hES cells in the presence of BMP/TGFβ inhibitors as described in Fig. 3A. Magnified regions corresponding to the insets are also shown. **(B)** Immunofluorescence analysis of the expression of ISL1/2 and NF-H in day 24 cultures differentiated from indicated hPSC lines in the presence of BMP/TGFβ inhibitors as described in Fig. 3A. Magnified regions corresponding to the insets are also shown. **(C)** Quantification of cells marked by expression of MN-associated proteins at D14 after differentiation of indicated MIFF1 hiPSCs in the presence of BMP/TGFβ inhibitors as described in Fig. 3A, and following immunofluorescence and image analysis. The data in the graph were obtained after scoring five random fields per experiment (two independent replicates). Error bars = s.d. **(D)** qPCR expression analysis of indicated thoracic, MN progenitor and MN markers at day 24 of differentiation of a hES (H9) and hiPS (SFCi55-ZsGr) cell line following freezing and thawing of cultures at D14 vs continuous differentiation controls.

**Figure S4.**
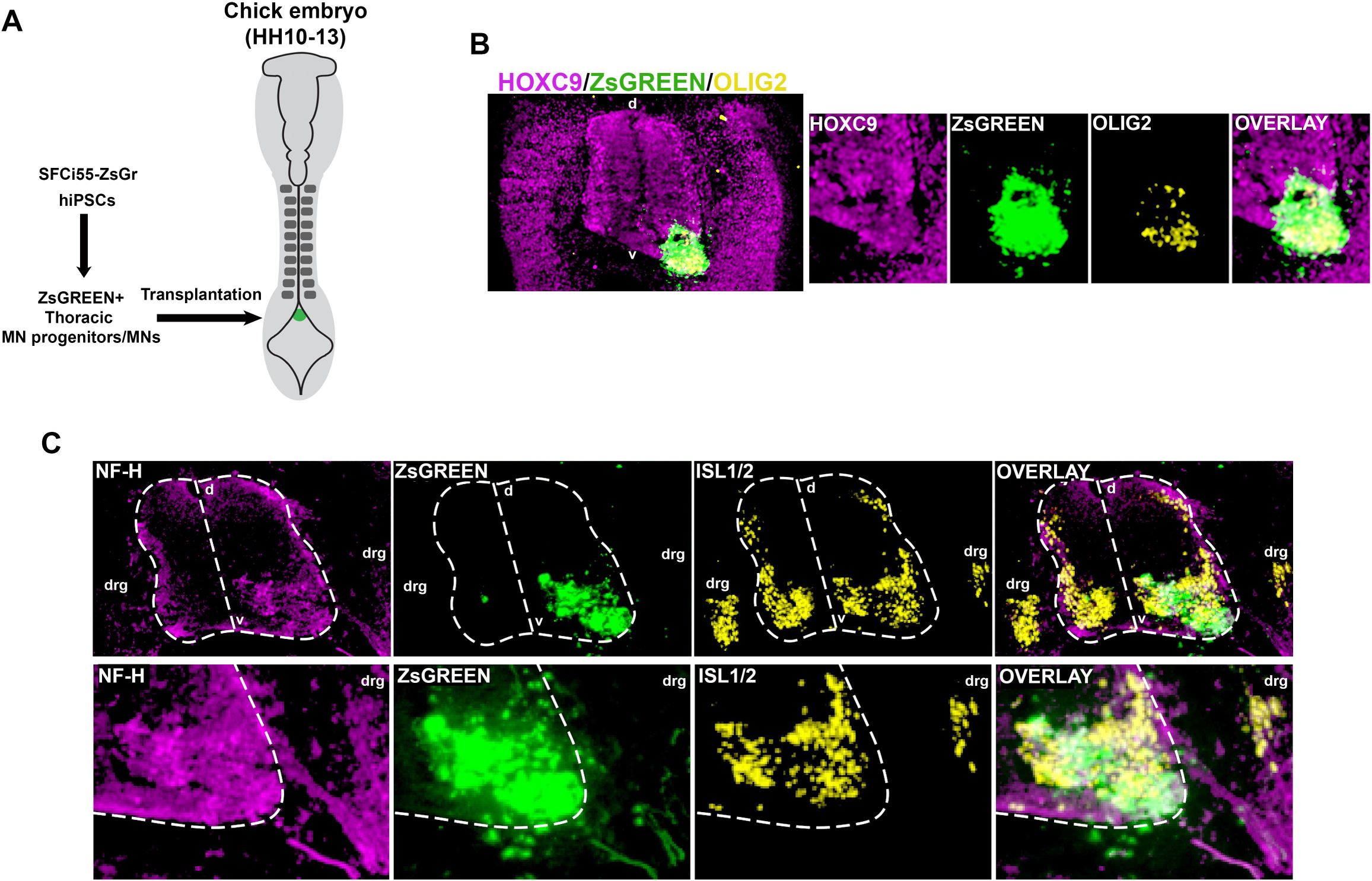
**(A)** Diagram depicting the *in vitro* generation of D14 posterior MN progenitors/MN from hPSCs and their transplantation into the posterior neural tube of HH10-13 chick embryos. **(B)** Immunofluorescence analysis of ZsGREEN, HOXC9 and OLIG2 expression in sections of HH26-30 chick embryos grafted with hPSC-derived ZsGREEN^+^ posterior D14 MN progenitor/MN cultures. Higher magnification images (right) indicate the co-expression of ZsGREEN and OLIG2 in a fraction of the grafted donor cells. Note that we employed an antibody that only detects the human version of OLIG2. D, dorsal; v, ventral. **(C)** Immunofluorescence analysis of ZsGREEN, NF-H and ISL1/2 expression marking donor human cells, neuronal projections and dorsal root ganglion (drg)/MN-containing ventral neural tube respectively, in sections of HH26-30 chick embryos grafted with hPSC-derived ZsGREEN^+^ posterior D14 MN progenitor/MN cultures. Higher magnification images (bottom row) indicate the co-expression of ZsGREEN and ISL1/2 in a fraction of the grafted donor cells. Dotted outlines mark the boundaries of the neural tube. D, dorsal; v, ventral.

**Figure S5.**
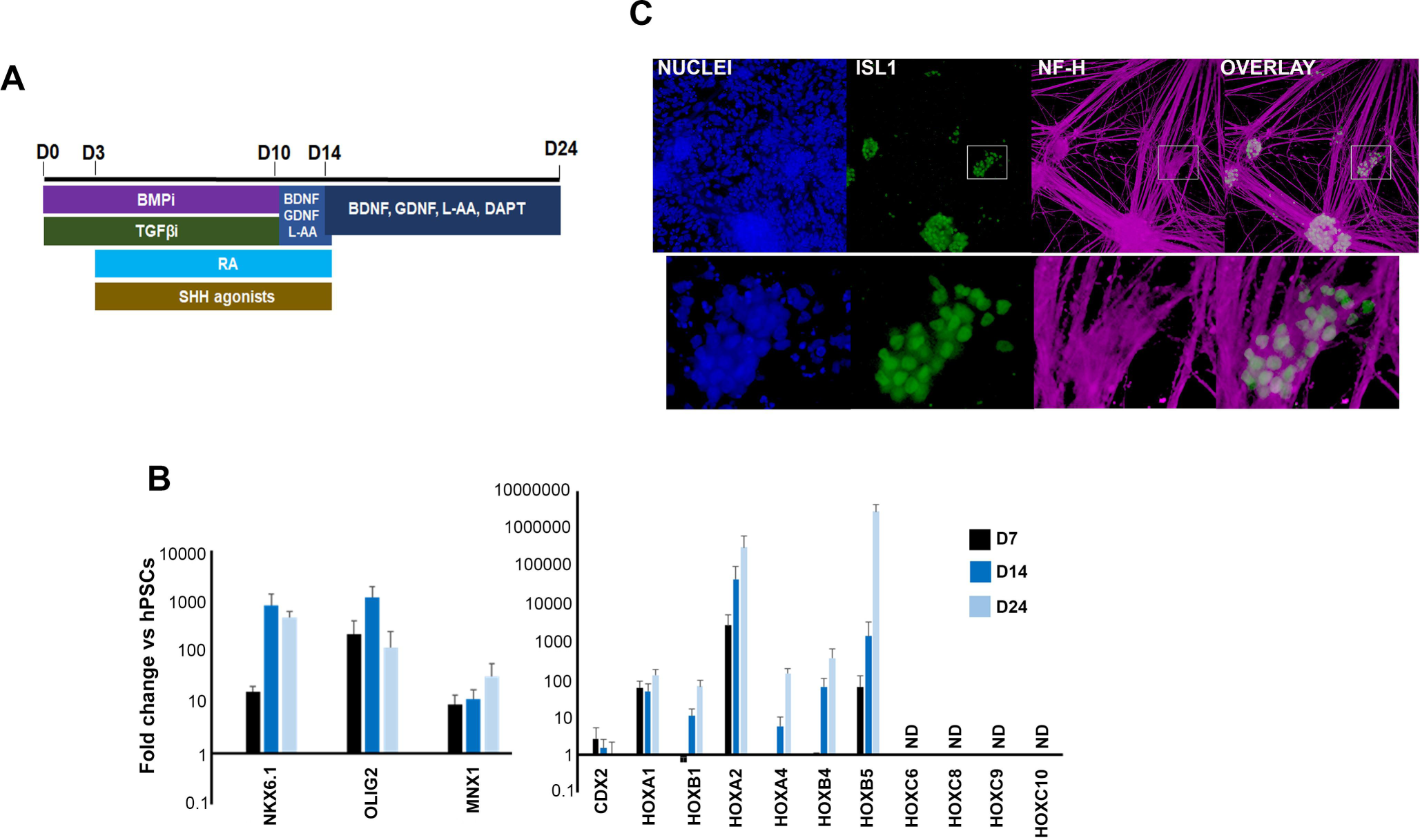
**(A)** Diagram depicting the culture conditions employed for the in vitro differentiation of hPSCs toward MNs of a hindbrain/cervical spinal cord identity. **(B)** qPCR expression analysis of indicated MN progenitor/MN markers (left) and HOX genes/CDX2 (left) at different points during the generation of MNs from hPSCs using the culture regimen shown in Fig. S5A. ND, Not detected. **(C)** Immunofluorescence analysis of the expression of ISL1/2 and NF-H in day 24 anterior MN cultures following differentiation of hPSCs as described in A. Magnified regions corresponding to the insets are also shown.

**Table S1.** Significantly up- and downregulated transcripts in day 7 early spinal cord progenitors vs day 3 NMP-like cells and part of GO enrichment analysis indicating upregulation of neural transcripts.

